# An integrated biobehavioral study of empathy

**DOI:** 10.1101/2025.11.27.690936

**Authors:** Luis Carretié, Fátima Álvarez, Uxía Fernández-Folgueiras, Oriana Figueroa, Estrella Veiga-Zarza, Miguel Pita

## Abstract

Empathy, a crucial prosocial human trait, is mainly measured through questionnaires, but performance in tasks involving emotional stimuli, brain activity during these tasks, and genetic information also provide relevant clues. These approaches are developed separately or by combining only some of these factors. Integrating them into a single study and analysis may help to better characterize their relative load and to understand the complexity of empathy. The IRI empathy scale was administered to 65 participants, who later performed a lexical decision task on emotional words while their performance and event-related potentials (ERPs) were recorded. Three genetic polymorphisms, one in the *OXTR* gene (SNP rs2254298) and two in the *AVPR1a* gene (RS1 and RS3 regions), were also analyzed. Generalized linear models revealed that these biobehavioral factors jointly predicted emotional but not cognitive empathy. Reaction times and the amplitude of the LPP component of ERPs in response to emotional words, along with rs2254298 AA/AG genotype, were significantly associated with scores on the IRI personal distress subscale. Meanwhile, LPP amplitudes and *AVPR1a*-RS1 LL genotype significantly predicted scores on the empathy concern subscale. The balanced loading of all these factors in the models reinforces the desirability of a comprehensive, multifaceted approach to characterizing empathy.

## 1. INTRODUCTION

Empathy is a key ingredient of human personality whose intensity largely determines social impulses and behaviors. In particular, it has been detected as a trait presenting high load in prosocial behavior (Davis, 2015; Decety et al., 2016) and low load in anti-social behavior such as bullying, callousness and psychopathic traits (Waller et al., 2020; Van Noorden et al., 2015). Empathy is currently conceptualized as a multifaceted rather than a monolithic trait. In general, there is a consensus in distinguishing at least a cognitive facet, consisting of being aware of others’ states, or being able to take the perspective of others, and an emotional one, consisting of being able to feel, to some extent, the emotions of others (Davis, 1983; Perry & Shamay-Tsoory, 2013; Zaki, 2017). These different experiential facets or layers of empathy are separately explored through certain empathy scales, such as the *Interpersonal Reactivity Index* (IRI; Davis, 1983).

However, indices of empathy besides scales have also been reported in different separate studies, some exploring neural activity and others measuring the performance in several tasks involving empathy to some extent. A frequent strategy in these studies is presenting emotional visual stimuli, such as facial expressions, emotional scenes, or affective words, given that empathy is assumed to modulate the processing of these stimuli (e.g., the processing of stimuli associated with others’ unpleasantness). On the one hand, studies on neural activity suggest that empathy modulates brain activity in response to these visual stimuli. Specifically, event-related potentials (ERPs) have revealed the effects of empathy at early and late processing stages by analyzing the relationship between the amplitude of different ERP components and scores in subscales measuring cognitive and affective facets of empathy. Results are complex -due in part to the heterogeneity of the experimental designs- and their exhaustive review exceeds our scopes but, in general, empathy -both cognitive and affective, although both facets are not always disentangled-modulates the brain response (for a recent systematic review see Almeida et al., 2024). Such modulation is more consistent and appears in response to a wider range of stimuli (scenes, faces and linguistic material) in late ERP positivities, such as the LPP component, than early latencies, which show modulation only to faces (Almeida et al., 2024). In this vein, the LPP evoked by emotional and neutral words, of particular interest here as explained below, has been reported to correlate with empathy scores (although not distinguishing between the cognitive and affective layers: Chou et al., 2020). On the other hand, studies on behavioral performance report better recognition -in terms of error rates-of emotional stimuli, particularly negative, as compared to neutral as empathy scores increase (without distinction between cognitive and affective components: Chikovani et al., 2015; faces). Relatedly, ambiguous expressions tend to be recognized as negative by empathic individuals (again, not distinguishing cognitive and affective facets: Fang & Li, 2024; faces). However, more recent studies reveal that emotion recognition may be modulated differently by each empathy facet as measured through IRI subscales (Israelashvili et al., 2020; faces).

Importantly, the level of empathy characterizing each individual seems to show stability throughout life (Knafo et al., 2008), suggesting, at least partially, a genetic regulation. The genetic profile may constitute, therefore, an additional marker of empathy. According to twin data, the heritability of empathy is more pronounced regarding the emotional than the cognitive component (Abramson et al., 2020). The most explored genes regarding empathy are those in charge of expressing vasopressin (AVP) and oxytocin (OXT) postsynaptic receptors in the brain. These two neuropeptides, with similar structure given that they evolved integratedly (Carter, 2017), may interact and even attach to mutual receptors. Both OXT and AVP neurons reach similar brain regions involved in social-emotional processes such as the amygdala, prefrontal areas, and the striatum, among others (Meyer-Lindenberg et al., 2011). In general, their effect, although complex, leads to diverse behaviors that may be conceptualized as prosocial (e.g., Donaldson & Young, 2008). Interestingly, several polymorphisms in *OXTR* and *AVPR1a* -the genes responsible for the expression of receptors for both neuropeptides-have been proposed to be associated with empathic behavior. Regarding *OXTR*, single nucleotide polymorphisms (SNP) such as rs2254298 present different allelic frequencies in high and low empathy individuals, particularly involving the presence or not of the A (for adenosine) allele (Christ et al., 2014). However, other *OXTR* SNPs have also been reported to be linked to empathy and prosocial behavior (e.g., Kohlhoff et al., 2022). With respect to AVP, the activation of V1aR receptors has been linked to empathy, with the *AVPR1a* gene being a target in several studies, which may present repeated sequences in some individuals (and hence, greater receptor expression). Particularly, sequence repeats in the RS3 region of this gene have been linked to non-aggressive behaviors (Vollebregt et al., 2021) and to high cognitive empathy scores (e.g., Uzefovsky et al., 2015). Some indirect clues also point to a link between the RS1 region of *AVPR1a* and empathy. Particularly, shorter alleles of RS1 may increase susceptibility to the autism phenotype (Tansey et al., 2011; Wassink et al., 2004), which is known to be characterized by low empathy scores and behaviors (e.g., Baron-Cohen et al., 2014).

The complexity of human traits, empathy being not an exception, makes a single marker (either subjective -scales-, behavioral, neural or genetic) insufficient to characterize them. This study aims to integrate and interrelate these four levels, separately explored up to the present. The scope is, first, to better characterize empathy nature, supporting or not the genetic influences mentioned above, but also confirming or not whether empathy leads to an integrated set of electrophysiological and performance markers that may help to detect and study this personality trait in the future. With this scope in mind, we carried out a study i) measuring separately the emotional and cognitive facts of empathy through the IRI scale, ii) determining the *OXTR* rs2254298 and *AVPR1a*-RS1 and -RS3 polymorphisms present in each participant (involved in OXT and AVP receptor expression), iii) measuring the performance in a lexical task involving emotional and neutral word processing, and iv), analyzing ERPs, and particularly the LPP component, during this same task. Analyses will introduce all these factors in a single generalized linear model to explore their mutual interrelationships and to define a hierarchy for their efficiency as empathy markers.

## 2. METHODS

### 2.1. Participants

Sixty-eight individuals participated in this experiment completing all phases, although data from only 65 of them could eventually be analyzed for data quality reasons explained below (38 women and 27 men, mean age=19.60, SD= 1.63). All participants were students of Psychology at the Universidad Autónoma de Madrid (UAM), voluntarily took part in the study, provided their informed consent according to the Declaration of Helsinki, and received academic compensation. The study had been approved by the UAM’s Ethics Committee (CEI 86-1626).

### 2.2. Empathy subjective measures

All participants completed a battery of personality scales. This battery also included scales to explore narcissism and psychopathy traits for purposes not relevant to the present study. As regards empathy, the battery included the IRI scale (Davis, 1983) which, as explained in the introduction, includes subscales reflecting cognitive empathy (perspective taking -IRIpt- and Fantasy -IRIf-), and subscales exploring emotional empathy (empathic concern -IRIec- and personal distress, IRIpd). These tests were administered in person using Google Forms, and the total time to fill out the entire battery was about 45 minutes on average.

The N=65 final sample was divided into two groups on each of the IRI subscales: the High group included participants with scores equal to or above the mean of the entire sample, while the Low group consisted of those with scores below the mean. Table 1 shows the gender distribution and descriptive statistics of both age and IRI subscales for High and Low groups. As may be appreciated, the resulting N for each group reached 29 participants in six subgroups out of eight, which our *a priori* sample size estimations yielded as necessary for a one-factor repeated-measures ANOVA (with three levels, as explained below) to reach a statistical power of 0.8^1^. Scores in High and Low groups were significantly different in all subscales (p<0.001 in all cases), whereas age did not differ between both groups in any subscale (p>0.09 in all cases; see Table 1 for details).

**Table 1.**
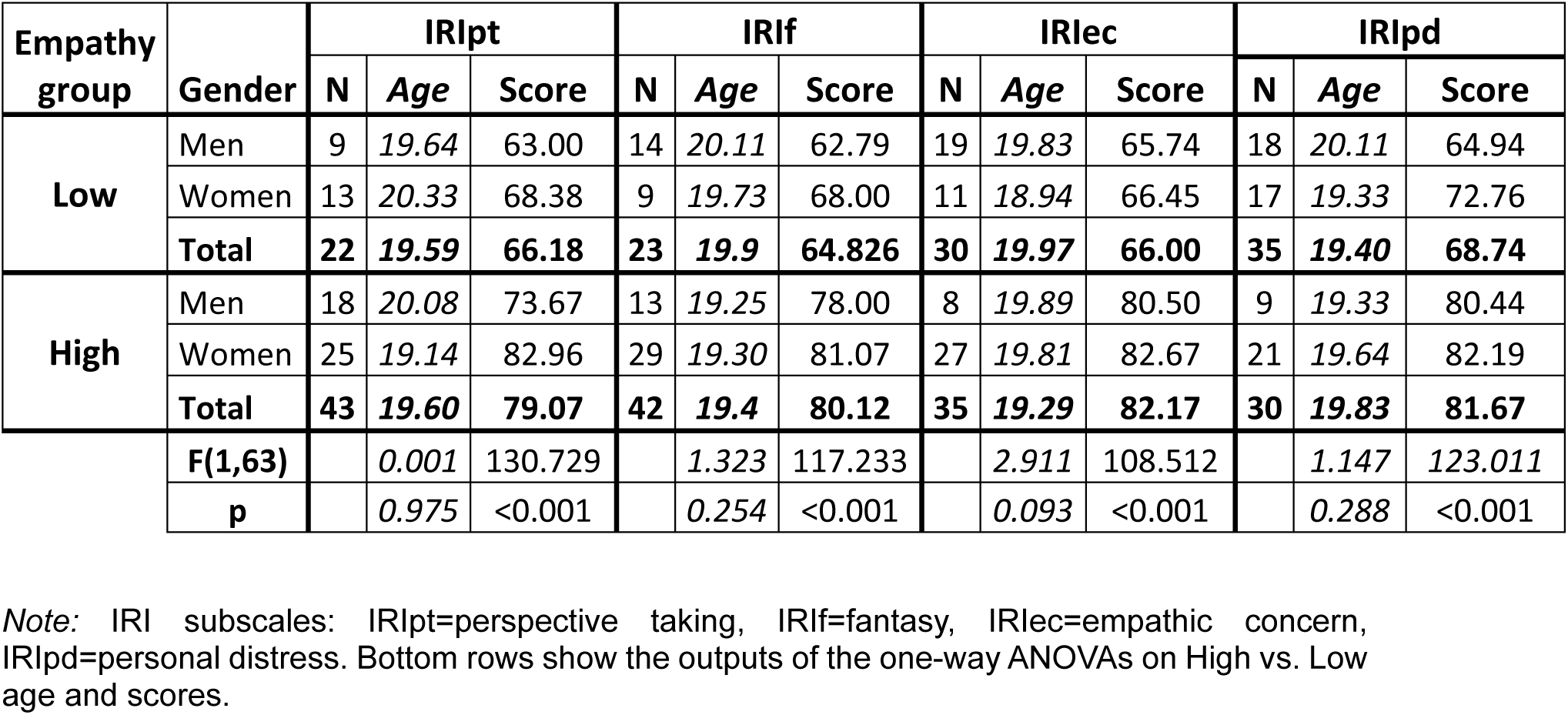
Distribution of men and women, mean age, and mean score, in the High and Low score groups within each IRI subscale.

| Empathy group | Gender | IRIpt |  |  | IRIf |  |  | IRlec |  |  | IRIpd |  |  |
| --- | --- | --- | --- | --- | --- | --- | --- | --- | --- | --- | --- | --- | --- |
|  |  | N | Age | Score | N | Age | Score | N | Age | Score | N | Age | Score |
| Low | Men | 9 | 19.64 | 63.00 | 14 | 20.11 | 62.79 | 19 | 19.83 | 65.74 | 18 | 20.11 | 64.94 |
|  | Women | 13 | 20.33 | 68.38 | 9 | 19.73 | 68.00 | 11 | 18.94 | 66.45 | 17 | 19.33 | 72.76 |
|  | **Total** | **22** | **19.59** | **66.18** | **23** | **19.9** | **64.826** | **30** | **19.97** | **66.00** | **35** | **19.40** | **68.74** |
| High | Men | 18 | 20.08 | 73.67 | 13 | 19.25 | 78.00 | 8 | 19.89 | 80.50 | 9 | 19.33 | 80.44 |
|  | Women | 25 | 19.14 | 82.96 | 29 | 19.30 | 81.07 | 27 | 19.81 | 82.67 | 21 | 19.64 | 82.19 |
|  | **Total** | **43** | **19.60** | **79.07** | **42** | **19.4** | **80.12** | **35** | **19.29** | **82.17** | **30** | **19.83** | **81.67** |
|  | **F(1,63)** |  | 0.001 | 130.729 |  | 1.323 | 117.233 |  | 2.911 | 108.512 |  | 1.147 | 123.011 |
|  | **p** |  | 0.975 | <0.001 |  | 0.254 | <0.001 |  | 0.093 | <0.001 |  | 0.288 | <0.001 |

*Note:* IRI subscales: IRIpt=perspective taking, IRIf=fantasy, IRlec=empathic concern, IRIpd=personal distress. Bottom rows show the outputs of the one-way ANOVAs on High vs. Low age and scores.

### 2.3. Lexical decision task

The second phase of the study consisted of a lexical decision task on emotional words during which participants’ behavioral and ERP activity were recorded.

#### 2.3.1. Stimuli and procedure

Participants were placed in an electrically shielded, sound-attenuated room. They were seated at a fixed distance (60 cm) from the screen (VIEWpixx®, 120 Hz) throughout this session. After placing the EEG cap (see “Recording and pre-processing” below), a sequence of words was presented on the screen. The list of words is available in Supplemental material, and consisted of 25 emotionally negative adjectives (e.g., “odioso”, -*hateful-*, “psicópata” -*psychopath*-), 25 positive (“chistoso” -*funny*-, “carismático” -*charismatic*-), and 25 neutral (e.g., “templado” -*temperate*-, “cartesiano” -*cartesian*-). We presented each adjective twice, so the number of trials was 150 (50 per emotional category), presented in random order. The selection was taken from the Hinojosa et al. (2016) adjective list, which includes normative data regarding emotional valence and arousal. Table 2 shows the average frequency of use of each adjective group, as well as their valence and arousal as reported in Hinojosa et al. (2016). An ANOVA was carried out to test whether the three categories differed in frequency of use, and confirmed no significant differences (F(2,72)=0.75, p=0.478, η^2^=0.020). Instead, ANOVAs on valence and arousal ratings showed significant differences between the three groups (F(2,72)=329, p<.001, η^2^=0.901 and F(2,72)=12.1, p<.001, η^2^=0.251, respectively).

**Table 2.** Mean and standard error of the mean (SEM) of frequency of use, valence and arousal of each group of adjectives (as reported in Hinojosa et al., 2016).

|  | Freq. of use |  | Valence |  | Arousal |  |
| --- | --- | --- | --- | --- | --- | --- |
|  | Mean | SEM | Mean | SEM | Mean | SEM |
| **Positive** | 1.876 | 0.191 | 7.272 | 0.139 | 5.614 | 0.254 |
| **Neutral** | 1.641 | 0.214 | 5.048 | 0.124 | 4.172 | 0.239 |
| **Negative** | 1.976 | 0.190 | 2.382 | 0.141 | 5.420 | 0.174 |

Participants were instructed to look permanently at the fixation dot appearing in the center of the screen and, whenever a word appeared (also at the center, replacing the fixation dot), to press one key if it had four or more syllables, and a different key if it had less than four (37 and 38 words -out of 75-, respectively). As illustrated in Figure 1, each word was presented during 400 ms, and the intertrial interval was 2500 ms. The adjectives were presented in Helvetica font and, as the fixation dot, were white on a grey background. The vertical visual angle of the highest letter case was 1.15°. The experimental run (150 trials) was divided into two blocks (75 trials each) with a rest period in between, and in total lasted 10 minutes approximately. Previously, and to let participants familiarize themselves with the task, a block of six practice trials (consisting of neutral adjectives, half under 4 syllables, different from those in the 75-word list) was presented. All parameters (timing and color) were identical to those of the experimental run. This practice block was repeated if the participant failed in more than one word.

**Fig. 1.**
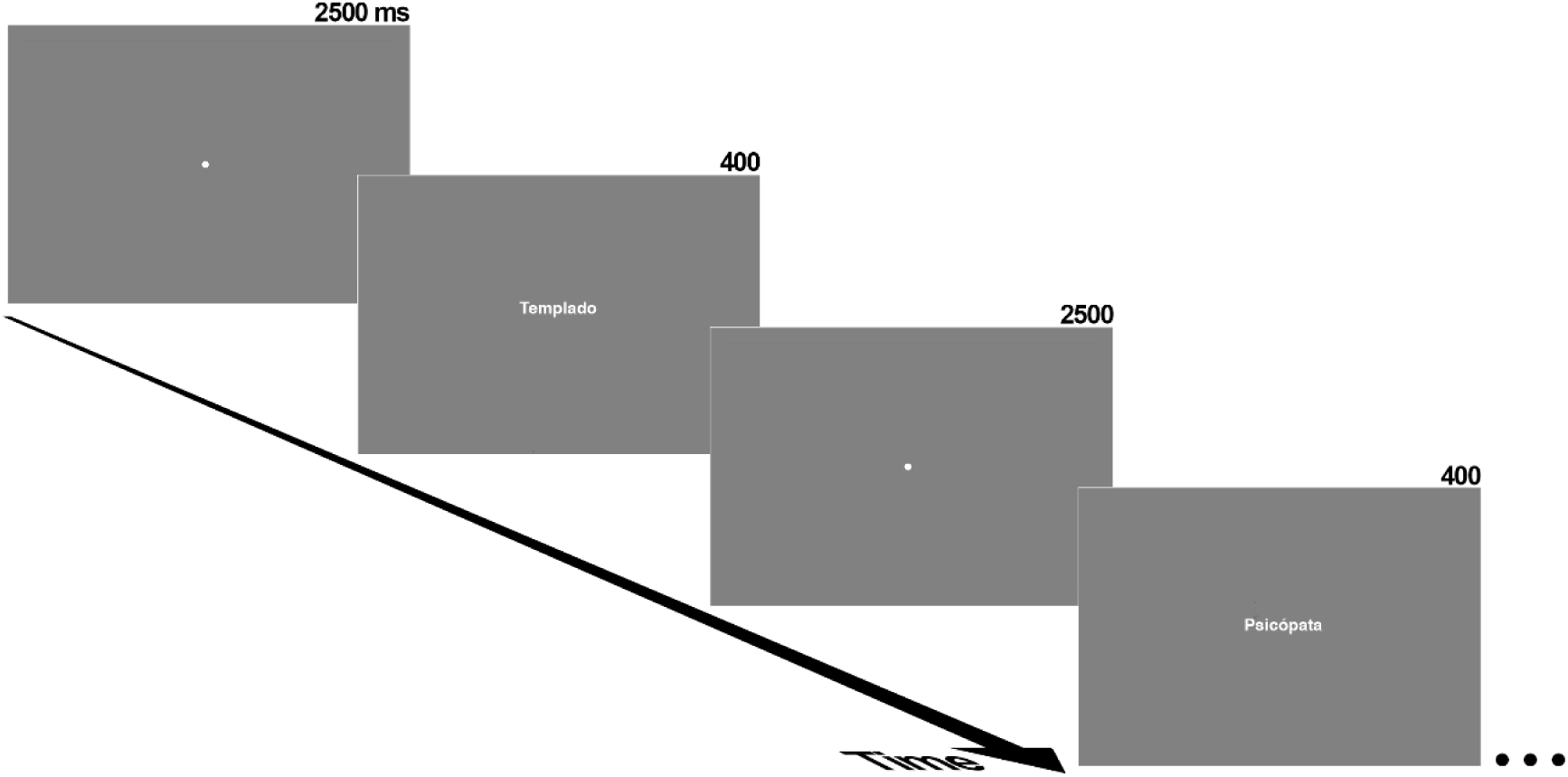
Schematic representation of one portion of the stimulus sequence. One example of neutral and emotionally negative adjectives is depicted.

#### 2.3.2. Recording and pre-processing

Electroencephalographic (EEG) activity was recorded using an electrode active cap (Biosemi) with Ag-AgCl electrodes. Sixty-four electrodes were placed at the scalp following a homogeneous distribution and the international 10-20 system. The EEG signal was pre-amplified at the electrode. Following the BioSemi design, the voltage at each active electrode was recorded with respect to a common mode sense (CMS) active electrode and a Driven Right Leg (DRL) passive electrode, replacing the ground electrode. All scalp electrodes were referenced offline to the nosetip. Electrooculographic (EOG) data were recorded supra- and infraorbitally (vertical EOG) as well as from the left versus right orbital rim (horizontal EOG) to detect blinking and ocular deviations from the fixation point. An online analog low-pass filter was set to 104 Hz (5^th^ order, CIC filter). Recordings were continuously digitized at a sampling rate of 512 Hz. An offline digital Butterworth bandpass filter of 0.3 to 30 Hz (4^th^ order, zero phase forward and reverse –twopass-filter) was applied to continuous (pre-epoched) data using the Fieldtrip software (http://fieldtrip.fcdonders.nl; Oostenveld et al. 2011). The continuous recording was divided into 1000 ms epochs for each trial, beginning 200 ms before the probe stimulus onset. The inevitable lag between the marks signaling stimuli onsets (or ‘triggers’) in EEG recordings and their actual onset on the screen was measured employing a photoelectric sensor as described in https://www.youtube.com/watch?v=0BPwcciq8u8, and corrected during pre-processing.

EEG epochs corresponding to trials in which participants responded erroneously or not responded in the task (see the previous section) were eliminated. Blinking-derived artifacts were removed through an independent component analysis (ICA)-based strategy (Jung et al. 2000), as provided in Fieldtrip. After the ICA-based removal process, a second stage of visual inspection of the EEG data was conducted to manually discard trials in which any further artifact, ocular (horizontal or vertical motion) or other type, was present. This automatic and manual rejection procedure led to the average admission of 41.25, 41.83, and 40.74 (respectively for positive, neutral and negative trials; SDs: 3.86, 4.52, 3.81). The minimum number of trials accepted for averaging was 31 trials per participant and condition (i.e., each category presented in each location). Data from one participant from the initial n=68 sample was eliminated since he/she did not meet this criterion. Two other participants were also discarded due to poor quality of the DNA sample (consequently, and as indicated, the final sample size was n=65).

Finally, detection and quantification of the LPP component of ERPs was carried out through a covariance-matrix-based temporal principal components analysis (tPCA), a strategy that has repeatedly been recommended for these purposes (e.g., Chapman et al., 2004; Dien, 2010). In brief, tPCA computes the covariance between all ERP time points, which tends to be high between those involved in the same component and low between those belonging to different components. Once quantified in temporal terms, LPP topography at the scalp level was decomposed into its main spatial regions via a spatial PCA (sPCA) performed on temporal factor scores. sPCA provides a reliable division of the scalp into different regions or spatial factors. Basically, each spatial factor is formed with the scalp points where recordings tend to covary. Temporal and spatial factor scores are the parameters in which temporal and spatial factors can be quantified, and are linearly related to amplitudes. The decision on the number of factors to select both in tPCA and sPCAs was based on the scree test (Cliff, 1987). Extracted factors were submitted to promax rotation in the case of tPCA and varimax for sPCA, as they have been defended as optimal in the time and space domain, respectively (Dien, 2010).

### 2.4. Genetic tests

Finally, the third phase consisted of saliva collection for genotyping. Just after the electrode cap was removed, and while still in the recording room, participants were instructed to self-collect approximately 1 ml of saliva in a Falcon tube using the passive drool method. For this step, participants were again left alone, and the procedure was repeated if the amount of saliva collected was insufficient. This sample tube was immediately identified with the anonymized participant code and stored at −80° C to preserve DNA until isolation.

DNA of each participant was isolated according to the instructions of the kit QIAmp DNA Mini kit (Qiagen). After isolation, DNA was measured and quality tested in a Nanodrop spectrophotometer. *OXTR* SNP rs2254298 and *AVPR1a* RS1 and RS3 genotypes were obtained for each participant after specific amplification by Polymerase Chain Reaction (PCR). The PCR reactions were performed in a final volume of 25 ul in a mixture of 100 ng of DNA, 2 mM MgCl2 (Bioline), 200 μM dNTPs (Biotools), 10 pmol of each primer, and 1.25 U of Taq Polymerase (Bioline); reactions were carried out in a thermal cycler (Techne TC-512). The PCR amplifications of the SNP rs2254298 of the gene *OXTR* were performed with the primers forward 5’-TGAAAGCAGAGGTTGTGTGGACAGG-3’ and reverse 5’-GAAGAAACTGGGGTGGGCGTT-3’. *AVPR1a* amplifications were performed with the primers forward 5’-VIC-AGGGACTGGTTCTACAATCTGC-3’ and reverse 5’- ACCTCTCAAGTTATGTTGGTGG-3’ for the RS1 regions, and 5’-FAM-TCCTGTAGAGATGTAAGTGC-3’ (forward) and 5’- TCTGGAAGAGACTTAGATGG −3’ (reverse) for the RS3 regions. In these two cases, the forward primer carried a fluorescent molecule in the 5’ region for its subsequent detection in capillary electrophoresis.

The reactions were performed as follows: initial denaturation for 5 min at 94 °C; followed by 35 cycles of: 45 s at 94 °C, 45 s at 62 °C (*OXTR*)/55 °C (*AVPR1a*), and 45 s at 72 °C; and a final extension of 10 min at 72 °C. To verify the integrity of the PCR products, 1% agarose gels were run in 1X TAE (Tris-acetate-EDTA) and revealed with 1X Red Gel (Biotium). Finally, *OXTR*-PCR products were sequenced by the Sanger method, while *AVPR1a*-PCR products were analyzed by capillary electrophoresis. In both cases, an ABI PRISM 3100 Genetic Analyzer (Applied Biosystems) was used to generate the files containing either the nucleotide sequence (.ab1 for *OXTR*) or the relative size of the amplified fragments (.fsa for *AVPR1a*).

*OXTR* results were analyzed using Bioedit Sequence Alignment Editor (Hall, 2011) to ascertain the presence of G and/or A nucleotides in the SNP rs2254298. To simplify analyses, participants were categorized according to their genotype into two groups: A-carrier (AA or GA genotypes) or GG (following Nishitani et al., 2017 and others). RS1 and RS3 polymorphisms of *AVPR1a* were analyzed employing Peak Scanner Software (Applied Biosystems) to estimate the length of the amplified fragment. For the RS1 region, repeats of nucleotides (GATA)n(GT)n are responsible of the allelic differences among individuals. Accordingly, alleles larger than 314 bp are considered Large (L) and the rest Small (S) (following Tansey et al., 2011). To simplify, participants were categorized in either L-carrier (genotypes LL y LS) or SS (Nishitani et al., 2017). Conversely, RS3 polymorphism -repeats (CT)n(GT)n-was categorized as follows: fragments larger than 328 bp are considered Large (L) and the rest Small (S). Participants are categorized in either S-carrier (genotypes SS y LS) or LL (Nishitani et al., 2017).

### 2.5. Data analysis

Analyses comprised two stages. Given that the lexical decision task was measured through several dependent variables (DVs), both behavioral and neural, the first stage was aimed at detecting which DVs were sensitive to the task (i.e., reflected significant differences between negative, neutral, and positive adjectives). To this aim, each of the spatial factors in which LPP was distributed throughout the scalp (four factors, as we will see in Results section), and each of the behavioral performance parameters (accuracy and reaction times), were submitted to a repeated-measures ANOVA on factor Emotion (three levels: Neg, Neu, Pos). Degrees of freedom of the F ratios were adjusted using the Greenhouse-Geisser (G-G) ε correction when necessary. The Bonferroni correction for multiple comparisons was applied in post-hoc tests. The scope of these first stage-analyses did not require dividing the sample into high and low empathy groups.

The second stage was exploring the interrelationship between empathy ratings as reflected in IRI subscales and i) the behavioral parameter(s) selected in the previous stage, ii) the LPP scalp region(s) selected in the previous stage, and ii) the genotype regarding *OXTR* (rs2254298), and *AVPR1a*-RS1 and *AVPR1a*-RS3 polymorphisms. To this aim, we applied a generalized linear model (GLM), specifically a logistic regression model with a Bernoulli family and logit link function to examine the relationship between being in the high or low group in each IRI subscale (outcome) and the behavioral, LPP and genotype factors (predictors) just described. Therefore, four GLM, one per IRI subscale scores, were performed. Prior to GLM, both behavioral and LPP factors were reduced by subtracting, within each individual, negative-neutral data (resulting in a single “negative relative to neutral” or NEG_RN_ factor), and positive-neutral data (POS_RN_).

## 3. RESULTS

Main results of genetic tests are first presented, followed by the main experimental effects. Regarding the latter, and as indicated in Methods section, a two-step analytical procedure was performed: i) detecting which dependent variables, both at the neural and the ERP level, were sensitive to the emotional content of adjectives in the lexical decision task, ii) examining the relationship between empathy scores and these dependent variables along with participants genotype in the three polymorphisms explored.

### 3.1. Genetic tests

Figure 2 illustrates the distribution of *OXTR* (rs2254298), *AVPR1a*-RS1 and *AVPR1a*-RS3 polymorphisms in the global sample. As may be appreciated, analyses of the SNP rs2254298 yielded 45 participants carrying the GG or double guanine genotype (27 women) and 20 carrying the adenosine or A allele -AG or AA genotype- (11 women). Regarding the *AVPR1a*-RS1 polymorphism, 26 participants carried double long alleles -LL genotype- (15 women) and 39 carried the short or S allele -SL or SS- (23 women). Finally, as for the *AVPR1a*-RS3 region, 29 individuals presented the SS genotype (18 women) and 36 carried the L variant -LS or LL- (20 women). Table 3 shows the distribution of the three polymorphisms in the high and low groups of each IRI subscale.

**Fig. 2.**
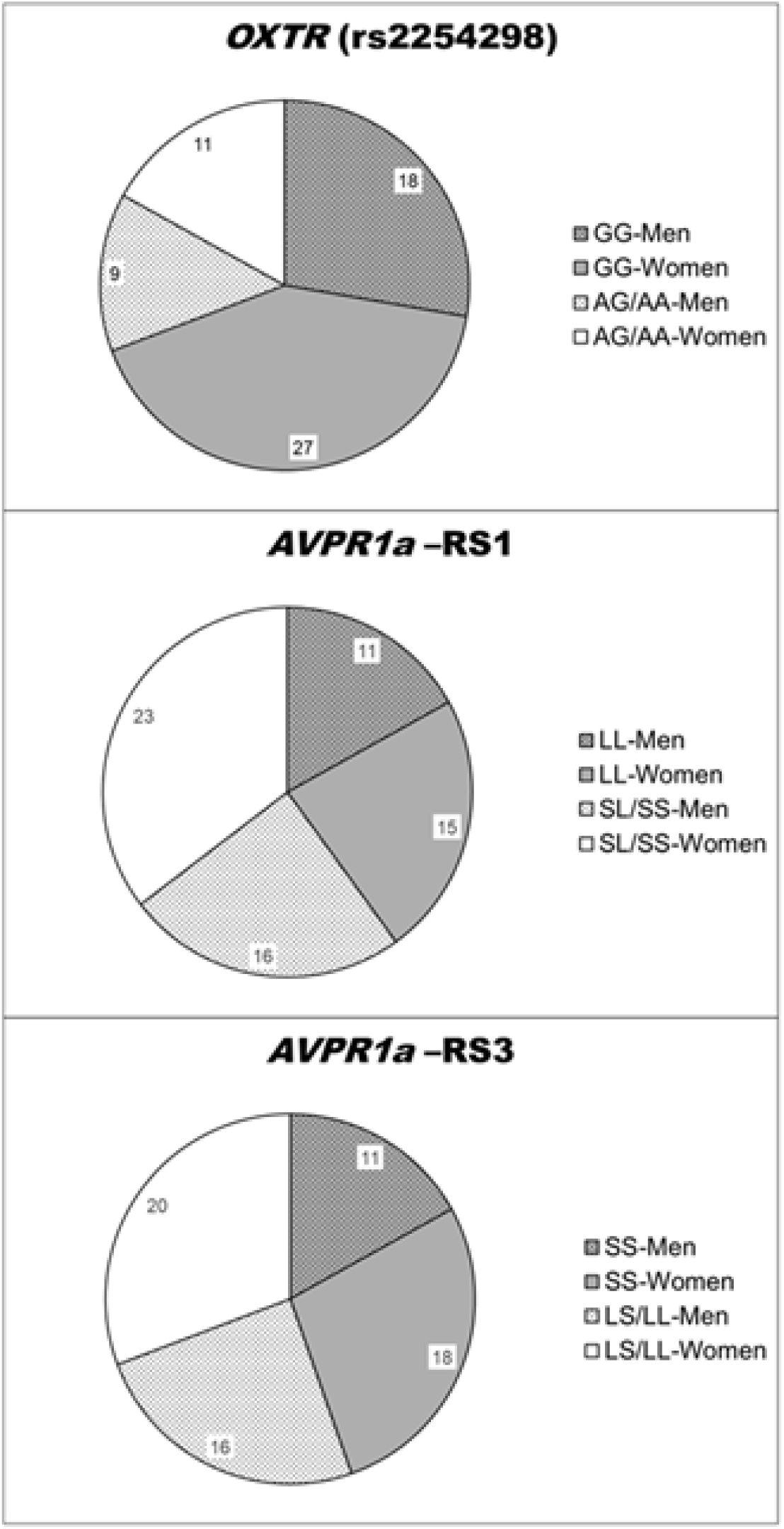
Allelic distribution of *OXTR* (rs2254298), *AVPR1a*-RS1, and *AVPR1a*-RS3 polymorphisms within the experimental sample.

**Table 3.** Allelic distribution within the high and low score groups for each IRI subscale.

| IRI subscale | Group | Allele | *OXTR*<br>(rs2254298) |  | *AVPR1a*-RS1 |  | *AVPR1a*-RS3 |  |
| --- | --- | --- | --- | --- | --- | --- | --- | --- |
|  |  |  | N | % | N | % | N | % |
| IRIpt | High | 1 | 17 | 37.78 | 10 | 34.48 | 7 | 26.92 |
|  |  | 2 | 5 | 25.00 | 12 | 33.33 | 15 | 38.46 |
|  | Low | 1 | 28 | 62.22 | 19 | 65.52 | 19 | 73.08 |
|  |  | 2 | 15 | 75.00 | 24 | 66.67 | 24 | 61.54 |
| IRIf | High | 1 | 14 | 31.11 | 11 | 37.93 | 7 | 26.92 |
|  |  | 2 | 9 | 45.00 | 12 | 33.33 | 16 | 41.03 |
|  | Low | 1 | 31 | 68.89 | 18 | 62.07 | 19 | 73.08 |
|  |  | 2 | 11 | 55.00 | 24 | 66.67 | 23 | 58.97 |
| IRIec | High | 1 | 20 | 44.44 | 12 | 41.38 | 6 | 23.08 |
|  |  | 2 | 10 | 50.00 | 18 | 50.00 | 24 | 61.54 |
|  | Low | 1 | 25 | 55.56 | 17 | 58.62 | 20 | 76.92 |
|  |  | 2 | 10 | 50.00 | 18 | 50.00 | 15 | 38.46 |
| IRIpd | High | 1 | 29 | 64.44 | 18 | 62.07 | 6 | 23.08 |
|  |  | 2 | 6 | 30.00 | 17 | 47.22 | 24 | 61.54 |
|  | Low | 1 | 16 | 35.56 | 11 | 37.93 | 20 | 76.92 |
|  |  | 2 | 14 | 70.00 | 19 | 52.78 | 15 | 38.46 |

*Note.* IRIpt=perspective taking, IRIf=fantasy, IRIec=empathic concern, IRIpd=personal distress. Both number of individuals (N) and percentages (%; referred to the total N within each allele) are provided. Allelic grouping for *OXTR* (rs2254298): 1=GG, 2=AG/AA; for *AVPR1a*-RS1: 1=SS, 2=LS/LL; for *AVPR1a*-RS3: 1=LL, 2=SL/SS.

### 3.2. Behavioral and neural sensitivity responses to emotional adjectives

Regarding performance in the lexical decision task, ANOVAs on reaction times revealed significant effects of factor Emotion (F(2,128)=17.172, G-G ε corrected p<0.001, η^2^=0.212). Post-hoc tests using the Bonferroni correction showed significantly faster responses to negative adjectives than to non-negative (both neutral and positive, p<0.001 in both cases; differences between these two categories were non-significant, p=1). The effects of Emotion on accuracy were also significant (F(2,128)=3.306, G-G ε corrected p=0.040, η^2^=0.049), but post-hoc tests failed to reach significance (all ps>0.079). Table 4 shows the main statistical descriptives of behavioral performance in the lexical decision task as a function of the emotional content of adjectives, and Figure 3 represents significant effects.

**Fig. 3.**
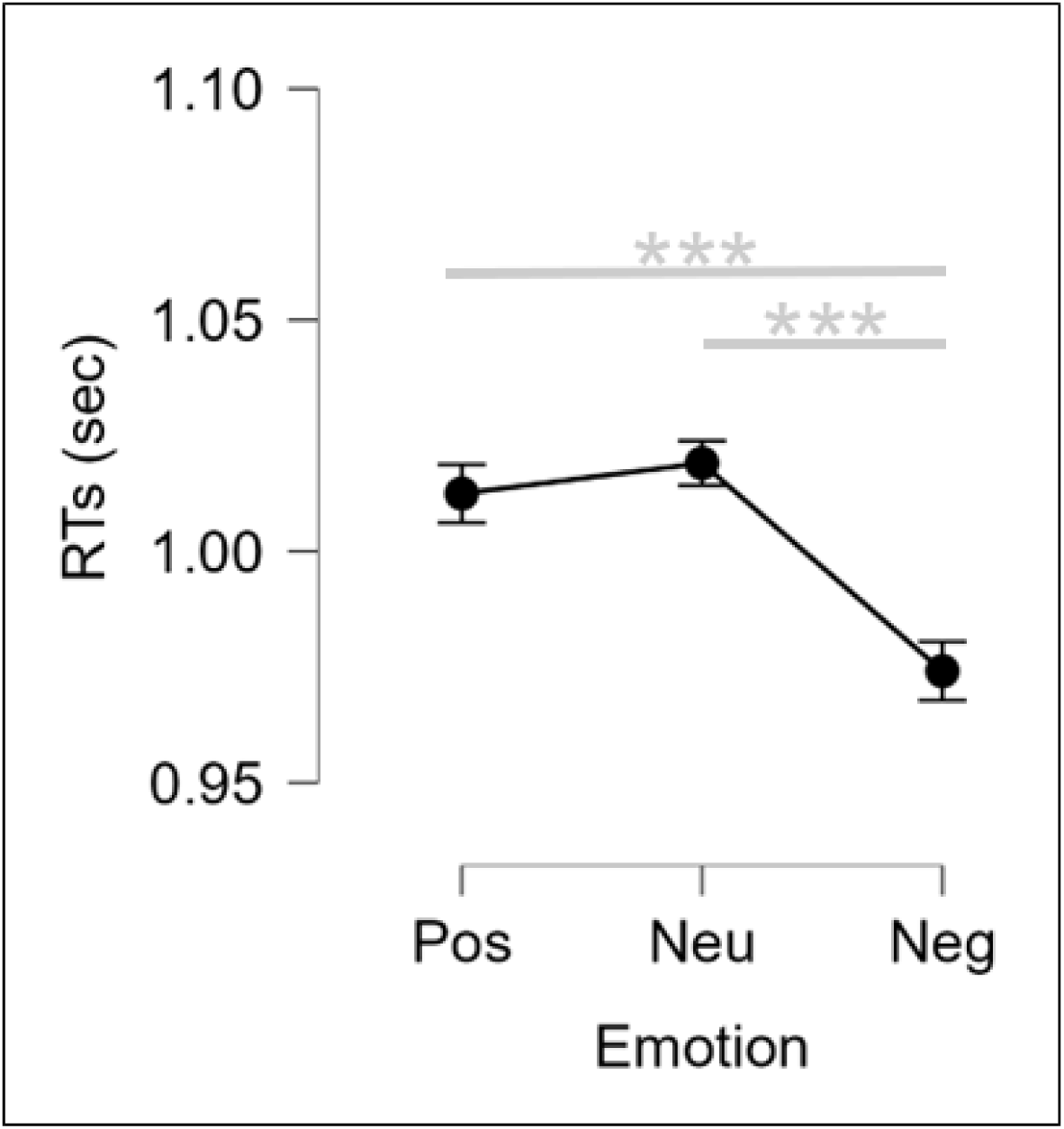
Means and standard error of means of reaction times (RTs) in response to the three types of adjectives: positive, neutral and negative (*** p<0.001)

**Table 4.** Means and standard error of means (SEM) of the two behavioral parameters (accuracy and reaction times -RTs-) and the five spatial factors of the LPP component of ERPs in response to adjectives as a function of their emotional load.

|  |  | Positive | Neutral | Negative |
| --- | --- | --- | --- | --- |
| Accuracy (0 to 1) | Mean | 0.883 | 0.898 | 0.883 |
|  | SEM | 0.008 | 0.009 | 0.008 |
| RTs (in seconds) | Mean | 1.013 | 1.019 | 0.974 |
|  | SEM | 0.041 | 0.039 | 0.036 |
| LPP-SF1 | Mean | -0.019 | 0.006 | 0.013 |
|  | SEM | 0.117 | 0.126 | 0.131 |
| LPP-SF2 (LPPa) | Mean | 0.118 | -0.168 | 0.05 |
|  | SEM | 0.129 | 0.122 | 0.121 |
| LPP-SF3 (LPPc) | Mean | 0.078 | -0.159 | 0.081 |
|  | SEM | 0.117 | 0.122 | 0.132 |
| LPP-SF4 | Mean | 0.035 | -0.038 | 0.003 |
|  | SEM | 0.125 | 0.117 | 0.132 |
| LPP-SF5 | Mean | -0.014 | -0.014 | 0.028 |
|  | SEM | 0.113 | 0.128 | 0.132 |

As for ERP responses in the task, Figure 4a shows a selection of grand averages after subtracting the baseline (prestimulus) activity. These grand averages correspond to medial prefrontal and central areas, where the experimental effects, discussed later, were most prominent. The tPCA (see Methods section) extracted six temporal factors (TFs). LPP, our target component as previously explained, was identified with TF6 given its latency and topographic distribution. The topography of FT6 was decomposed into five spatial factors (SFs) by sPCA. Figures 4b and 4c summarize, respectively, the tPCA and sPCA outputs. Subsequently, spatial factor scores corresponding to each SF (linearly related to amplitudes, as indicated) were submitted to five repeated-measures ANOVAs on factor Emotion (one ANOVA per spatial factor). Two of the LPP spatial factors, SF2 and SF3 (hereafter LPPa -anterior- and LPPc -central-to make results easier to understand) showed significant effects by the emotional content of adjectives (LPPa: F(2,128)=3.861, G-G ε corrected p=0.031, η^2^=0.057; LPPc: F(2,128)=6.864, G-G ε corrected p=0.002, η^2^=0.097): Figure 4d. Post-hoc tests with Bonferroni correction for multiple comparisons showed that LPPa amplitudes for positive adjectives were greater than for neutral (p=0.026); negative vs. neutral and positive vs. negative comparisons did not reach significance (p=0.135 and 1.000, respectively). Post-hoc tests regarding LPPc revealed greater amplitudes to both negative and positive adjectives than to neutral (p=0.005 in both cases), the difference between negative and positive being non-significant (p=1.000).

**Figure 4.**
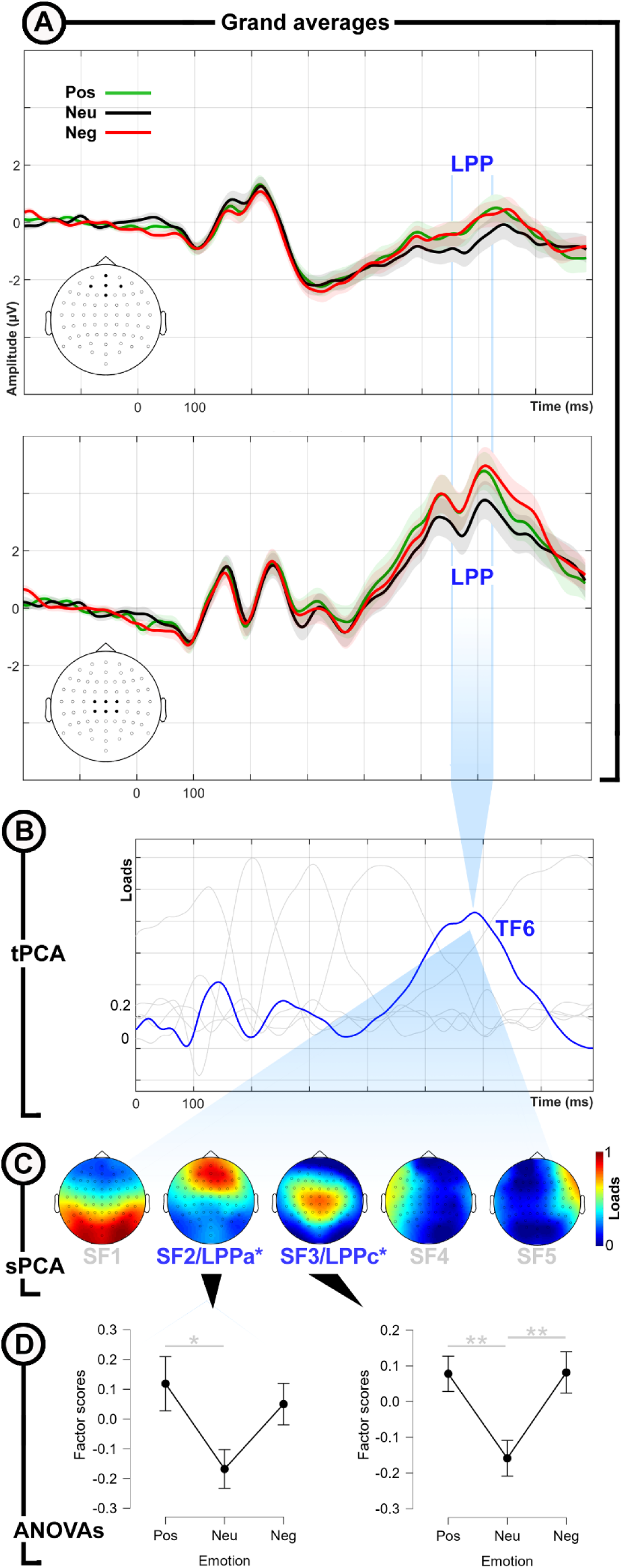
**A.** Grand averages of ERPs recorded at midline anterior and central scalp electrodes (marked in the schematic white heads), where the emotional word task produces significant effects. Shadows surrounding grand average lines represent the standard error of means. **B.** Temporal factors extracted through temporal principal component analysis (tPCA). TF6, which corresponds to LPP, is marked in red. **C.** Spatial factors in which LPP may be decomposed, according to spatial PCA (sPCA). Those showing significant effects in ANOVAs, LPPa and LPPc (see D), are labelled in blue fonts. **D.** Means and standard error of means of LPPa and LPPc amplitudes in response to the three types of adjectives: positive, neutral and negative (* p<0.05, ** p<0.01)

### 3.3. Multivariate relationships

As indicated, a GLM (Bernouilly family, logit link) was applied to each of the four IRI subscale groups (High and Low) in order to test whether, and how, RTs, LPPa and LPPc (the factors revealed as sensitive to the emotional content of adjectives in the lexical decision task; see previous section), as well as polymorphisms of *OXTR* (rs2254298), *AVPR1a*-RS1 and *AVPR1a*-RS3, predict together each empathy facet. Table 5 shows the main outputs of these analyses, and Figures 5 and 6 illustrate those with significant coefficients. Two of the subscales, IRIpd and IRIec, showed significant associations with the predictors.

**Fig. 5.**
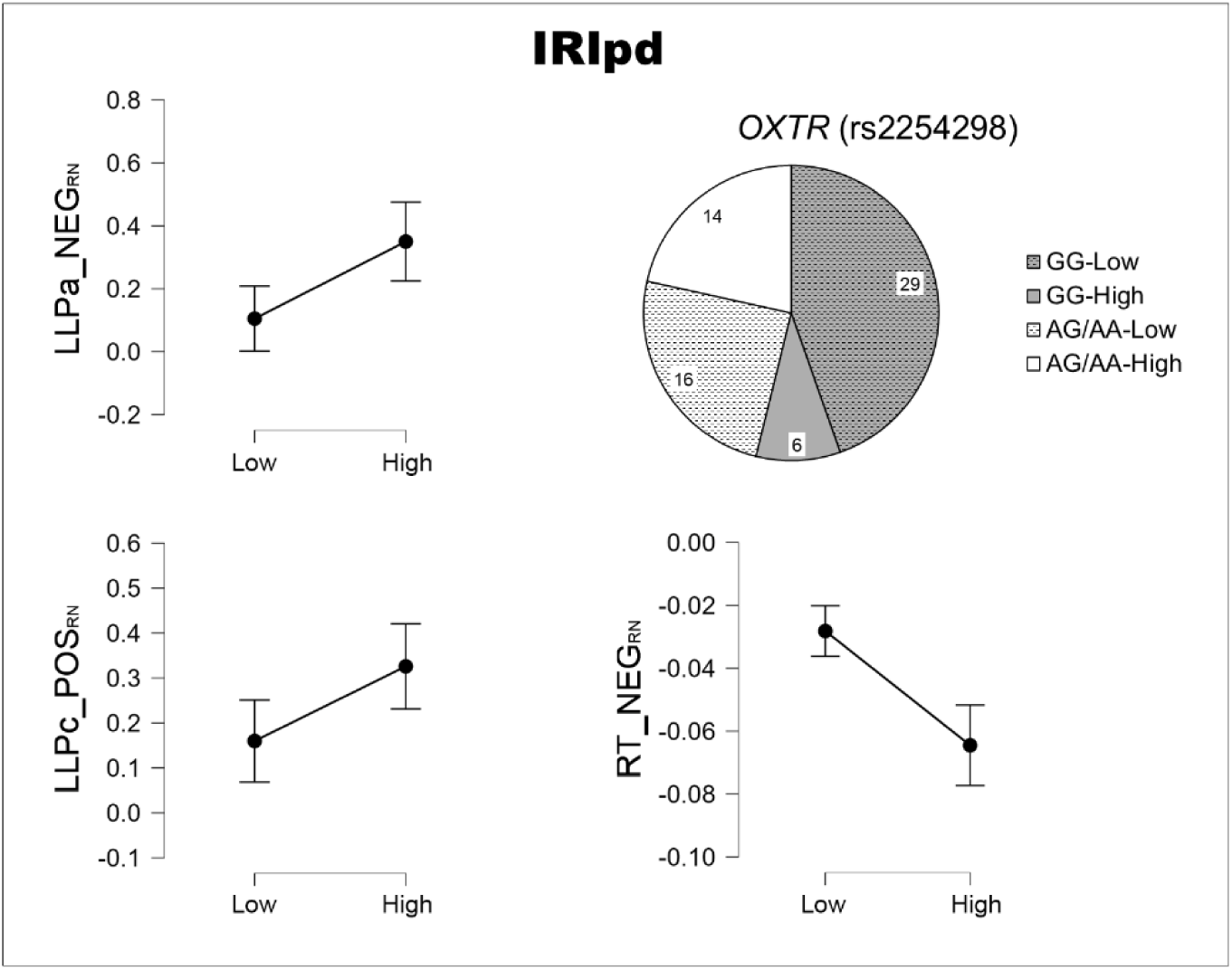
Coefficients corresponding to the individual predictors introduced in the generalized linear model on the IRIpd (personal distress) subscale. RTs=reaction times. The RN subindex in the predictor names means “relative to neutral” (i.e., subtracting the neutral from each emotional condition).

**Fig. 6.**
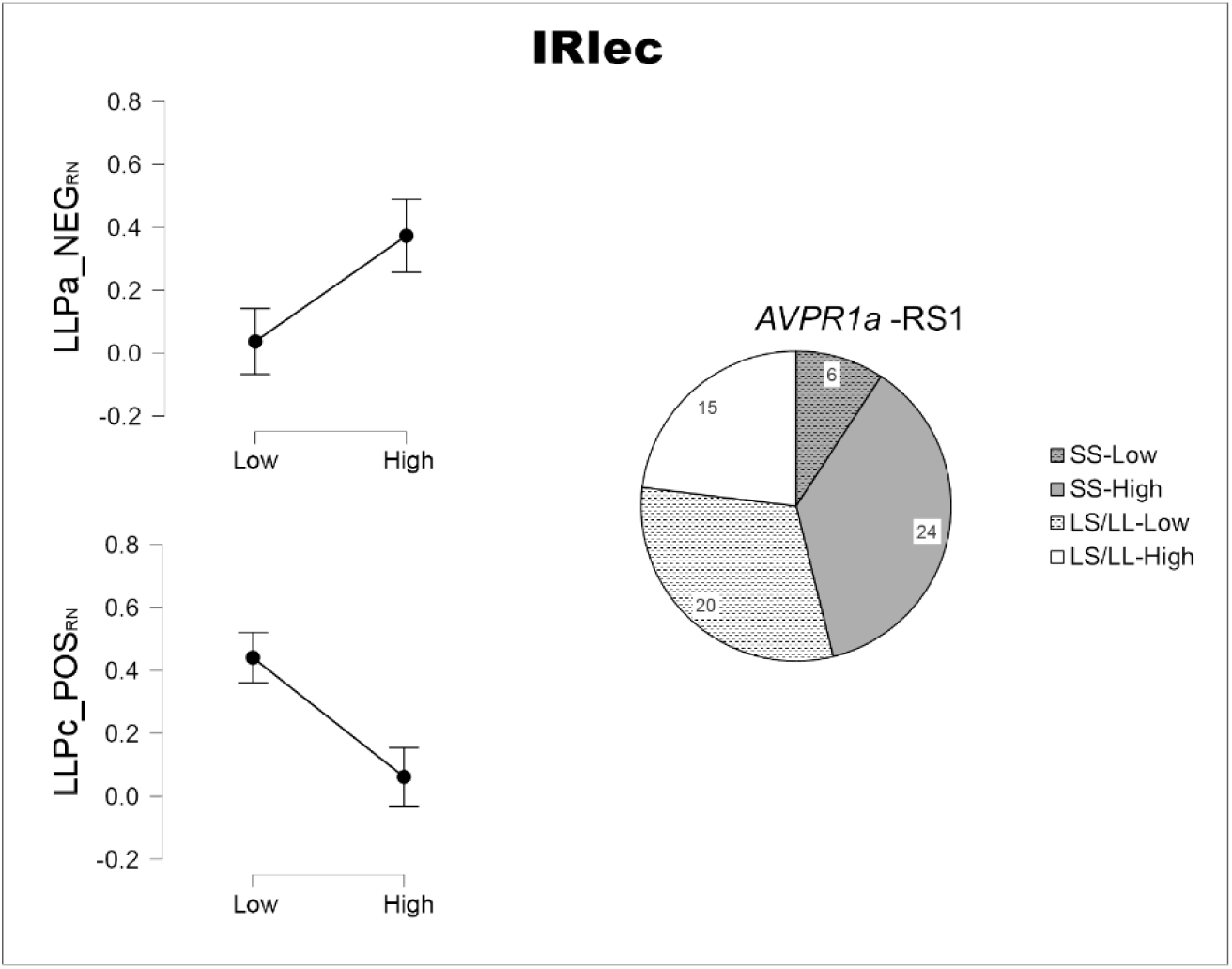
Coefficients corresponding to the individual predictors introduced in the generalized linear model on the IRIec (empathic concern) subscale. RTs=reaction times. The RN subindex in the predictor names means “relative to neutral” (i.e., subtracting the neutral from each emotional condition).

**Table 5.** Coefficients of individual predictors in the two GLMs where the models showed a significantly better fit than the null model: those on IRIpd (personal distress subscale) and IRIec (empathic concern) as the output.

|  | Predictors | Estimate | Standard Error | z | p |
| --- | --- | --- | --- | --- | --- |
| IRlpd | RTs-POS<sub>RN</sub> | -2.100 | 5.866 | -0.358 | 0.720 |
|  | RTs-NEG<sub>RN</sub> * | -18.857 | 6.786 | -2.779 | 0.005 |
|  | LPPa-POS<sub>RN</sub> | -0.771 | 0.458 | -1.681 | 0.093 |
|  | LPPa-NEG<sub>RN</sub> * | 1.622 | 0.670 | 2.422 | 0.015 |
|  | LPPc-POS<sub>RN</sub> * | 1.907 | 0.841 | 2.266 | 0.023 |
|  | LPPc-NEG<sub>RN</sub> | -0.996 | 0.662 | -1.504 | 0.133 |
|  | *OXTR* (rs2254298) * | 2.798 | 0.873 | 3.205 | 0.001 |
|  | *AVPR1a*-RS1 | -0.390 | 0.713 | -0.547 | 0.584 |
|  | *AVPR1a*-RS3 | 0.988 | 0.680 | 1.453 | 0.146 |
| IRlec | RTs-POS<sub>RN</sub> | -0.903 | 5.800 | -0.156 | 0.876 |
|  | RTs-NEG<sub>RN</sub> | -8.120 | 5.587 | -1.453 | 0.146 |
|  | LPPa-POS<sub>RN</sub> | -0.512 | 0.366 | -1.399 | 0.162 |
|  | LPPa-NEG<sub>RN</sub> * | 1.388 | 0.640 | 2.167 | 0.030 |
|  | LPPc-POS<sub>RN</sub> | -2.613 | 0.982 | -2.661 | 0.008 |
|  | LPPc-NEG<sub>RN</sub> * | -0.595 | 0.698 | -0.853 | 0.394 |
|  | *OXTR* (rs2254298) | -0.267 | 0.731 | -0.364 | 0.716 |
|  | *AVPR1a*-RS1 * | -2.579 | 0.826 | -3.123 | 0.002 |
|  | *AVPR1a*-RS3 | -0.235 | 0.697 | -0.337 | 0.736 |

\* Significant predictor
*Note.* RTs=reaction times. The RN subindex in the predictor names means “relative to neutral” (i.e., subtracting the neutral from each emotional condition). The asterisk marks a significant association between the individual predictor and the output ( $p < 0.05$ ).

Regarding IRIpd, the likelihood ratio test confirmed that the model provided a better fit than the null model (χ²(55) = 30.937, p <0.001). Deviance goodness-of-fit test for model fitness did not indicate any inadequacies (p = 0.553). As may be appreciated in Table 5, where coefficients for each predictor are detailed, reaction times to NEG_RN_ (RN means “relative to neutral” or “minus neutral”, as indicated in Methods), LPPa to NEG_RN_, LPPc to POS_RN_, and OXTR, were significant predictors of the IRIpd group to which participants belonged. Concretely, high IRIpd individuals were characterized by shorter RTs and greater LPPa amplitudes to NEG_RN_ than low IRIpd participants. Moreover, high IRIpd participants also showed greater LPPc amplitudes to POS_RN_ than low IRIpd. Concerning genes, the proportion of A-carriers in the rs2254298 polymorphism of the *OXTR* gene was greater in the high IRIpd group (70% of total A-carriers vs. 30%), whereas participants with genotype GG dominated the low IRIpd group (64.44% vs. 35.56%). The rest of predictors were not significantly associated with the outcome (p >0.090 in all cases, see Table 5).

As for IRIec, the likelihood ratio test confirmed that the model provided a better fit than the null model (χ²(55) = 31.905, p <0.001). Deviance goodness-of-fit test showed again a good fit of the model of interest (p = 0.372). Concerning coefficients (Table 6), LPPa to NEG_RN_, LPPc to POS_RN_, and RS1, were significant predictors of the participants’ IRIec group. In particular, high IRIec participants presented greater LPPa amplitudes to NEG_RN_ than low IRIec participants. Instead, low IRIec participants showed greater LPPc amplitudes to POS_RN_ than high IRIec ones. With respect to polymorphisms, the proportion of LL-carriers in the RS1 region of the *AVPR1a* gene was greater in the high IRIec group (76.92% of total LL carriers vs. 23.08%), while the proportion of S-carriers was greater in the low IRIec group (61.54% vs. 38.46%). The rest of predictors, including RTs, were not significantly associated with the outcome (p >0.146 in all cases, see Table 5).

## 4. DISCUSSION

Our study explored, in an integrated fashion, the behavioral, neural and genetic profiles characterizing empathy. To assess behavioral and neural responses, we presented participants with emotional words, to which they were asked to pay attention and make a lexical decision. We found that reaction times (in the behavioral domain) and both LPPa and LPPc (in the neural domain), were sensitive to the emotional content of the stimuli. As for the genetic profile, we analyzed three polymorphisms, one linked to oxytocin receptor expression in the brain -*OXTR* gene-, namely rs2254298, and two polymorphisms linked to a vasopressin receptor gene -*AVPR1a*-, particularly the sequence repeats in regions RS1 and RS3. Using generalized linear models, we explored the extent to which these factors, introduced together in the model, predicted the empathy scores in the cognitive and affective subscales of the IRI empathy test (Davis, 1983). Importantly, a key basic result is that behavioral, neural and genetic factors significantly predict emotional empathy (i.e., IRIpd and IRIec scores), but not cognitive empathy (IRIpt and IRIf). Next, a detailed discussion of the main results is presented.

Beginning with *personal distress*, and at the neural level, the amplitudes of LPPa and LPPc were greater in response to both negative and positive words, respectively, in the high IRIpd than in the low emotional empathy groups. This result is in line with previous studies also showing greater LPP amplitudes to positive and negative words in high-empathy individuals. The LPP has been consistently reported to reflect the interplay of emotional and attentional processes, showing enhanced amplitudes to emotional stimuli than to neutral, but also to attended emotional stimuli than to unattended ones (see, e.g., Hajcack et al., 2009; MacNamara et al., 2009). In terms of the two main affective dimensions, valence and arousal (Lang et al., 1993; Osgood et al., 1957; Russell, 1979), which range from calming to arousing and from negative to positive, respectively, high personal distress seems, therefore, to be characterized by an *attentional arousal bias.* Indeed, arousal is orthogonal to valence so both positive and negative stimuli may be arousing. However, these ERP results are qualified by behavioral performance data, which point to a relatively greater load for negative stimuli. Thus, reaction times to negative words, but not to positive words (relative to neutral in both cases), showed a significant association with IRIpd, consisting of shorter latencies in the high group. This result converges with previous data reporting better performance for negative visual stimuli in high-empathy individuals (not distinguishing cognitive from emotional empathy: Chikovani et al., 2015), and may reflect post-attentional processes, probably of lexical nature considering our task. Therefore, our neural and behavioral data point to personal distress to show increased attentional resources towards emotional stimuli in general, but an advantage towards negative items in post-attentional stages, either cognitive and/or underlying the final motor output.

Contrarily, a sort of *attentional valence bias* was observed regarding *empathic concern*. Indeed, LPPa and LPPc amplitudes were higher to negative stimuli in the high IRIec group and to positive stimuli in the low IRIec group, respectively. As indicated, LPP may be reflecting enhanced attentional mechanisms, in this case in different valence directions in the high and low groups. The attentional bias towards negative stimuli in high empathic concern individuals is an expected result given their drive and capability to “read” others’ emotional states (Batson et al., 1987; Davis, 1983), but the explanation for the “positivity bias” in the low emotional empathy group as compared to the high group is not straightforward. In our opinion, it may reflect a sort of “negativity inhibition” achieved either through attentional amplification towards positive items in detriment of negative, or through active attentional suppression, at least partial, of negative information (or through both mechanisms). Additional research is needed to further characterize this sort of “push and pull” attentional mechanism regarding emotional valence as a function of empathy.

Genes controlling the expression of oxytocin and vasopressin receptors in the brain have also contributed to the joint model to predict emotional empathy scores. This is in line with twin data which signal that the heritability of empathy affects its emotional facet rather than the cognitive facet (Abramson et al., 2020). On one hand, the rs2254298 polymorphism of the *OXTR* gene significantly contributed to predicting IRIpd. Concretely, the proportion of the adenine allele carriers is much greater in the high than in the low IRIpd score group (70% vs. 30%). This result converges with previous studies also reporting greater IRIpd scores in individuals presenting the rs2254298 AA/AG genotype than those with the GG genotype (Li et al., 2023; this effect interacted with the s13316193 polymorphism). Indeed, the rs2254298 GG genotype has been associated with lower oxytocin plasma levels than AA/AG (Feldman et al., 2012). Crucially, lower oxytocin plasma levels have been linked to low levels of empathy (Barraza & Zak, 2009). On the other hand, analyses regarding the *AVPR1a* genotypes showed two relevant findings. First, repetitions in the RS3 region were not associated with any empathy subscale in our models, contrary to previous data reporting this association, particularly with cognitive empathy (Uzefovsky et al., 2015). This discrepant result could be due to the potentially low statistical power in our analyses on IRI cognitive subscales (IRIpt and IRIf), given that one of the groups (the low empathy one, in both cases), did not reach the desirable sample size (n<29). Second, repetitions in the RS1 region were strongly linked to greater empathy scores in the IRIec subscale, a novel result not reported previously to the best of our knowledge. However, this result is not surprising given that the shorter alleles of RS1 may be more prone to suffer autism (Tansey et al., 2011; Wassink et al., 2004), which at the same time is characterized by low empathy scores (Baron-Cohen et al., 2014). In any case, this result points to the relevance of this particular polymorphism in future research on the genetic correlates of empathy.

In sum, behavioral and neural responses to emotional stimuli, as well as certain *OXTR* and *AVPR1a* genotypes, predict emotional empathy. On one hand, reaction times and the amplitude of the LPP component of ERPs in response to emotional words, together with rs2254298 AA/AG genotype were significantly associated with the scores in the personal distress subscale of IRI. On the other hand, both LPP amplitudes and the *AVPR1a*-RS1 L-carriers significantly predicted scores on the empathy concern subscale. Whereas all these factors present a relatively balanced load in the model as regards their significance as predictors, which supports the convenience of a comprehensive and multifaceted strategy to study empathy, the LPP component of ERPs remains the most consistent index across emotional empathy subscales. Thus, while performance and each particular genotype (strongly) predict the scores in one of the emotional empathy subscales, LPP is “transversal” to both subscales. This points to this component as a particularly valuable index to exploring empathy. Future studies following a multifaceted biobehavioral strategy are needed to further characterize empathy. They would benefit from larger sample sizes to explore interactions such as gender x genotype, given the male vs. female differences in empathy scores as a function of their allele characteristics in the rs2254298 polymorphism (Christ et al., 2014). This integrated biobehavioral view could also be pursued by exploring other candidate polymorphisms that were not analyzed in our study, but also potentially relevant (e.g., SNP rs53576 of *OXTR* or the (GT)_25_ repeat of *AVPR1a*). Finally, including a wider variety of participants (e.g., increasing age variability and recruiting population samples characterized by more polarized empathy traits) would also be of interest in this multifactorial strategy.

## DATA AND CODE AVAILABILITY

The data associated with this experiment are available in our OSF repository (https://osf.io/z3reu/). Supplemental material mentioned hereafter is also available at that link.

## AUTHOR CONTRIBUTIONS

**Luis Carretié:** Conceptualization, Formal Analysis, Funding acquisition, Project administration, Supervision, Visualization, Writing – original draft. **Fátima Álvarez:** Data curation, Investigation, Project administration, Supervision, Writing – review & editing. **Uxía Fernández-Folgueiras:** Data curation, Investigation, Software, Writing – review & editing. **Estrella Veiga-Zarza:** Data curation, Investigation, Writing – review & editing. **Oriana Figueroa:** Methodology, Resources, Writing – review & editing. **Miguel Pita:** Conceptualization, Investigation, Resources, Writing – review & editing.

## Footnotes

1 We used MorePower 6.0.4 applet (Campbell & Thompson, 2012) introducing a medium effect size (eta^2^=0.08).

